# The Tradeoffs Between Persistence and Mutation Rates at Sub-Inhibitory Antibiotic Concentrations in *Staphylococcus aureus*

**DOI:** 10.1101/2024.04.01.587561

**Authors:** Alysha S. Ismail, Brandon A. Berryhill, Teresa Gil-Gil, Joshua A. Manuel, Andrew P. Smith, Fernando Baquero, Bruce R. Levin

## Abstract

The rational design of the antibiotic treatment of bacterial infections employs these drugs to reach concentrations that exceed the minimum needed to prevent the replication of the target bacteria. However, within a treated patient, spatial and physiological heterogeneity promotes antibiotic gradients such that the concentration of antibiotics at specific sites is below the minimum needed to inhibit bacterial growth. Here, we investigate the effects of sub-inhibitory antibiotic concentrations on three parameters central to bacterial infection and the success of antibiotic treatment, using *in vitro* experiments with *Staphylococcus aureus* and mathematical-computer simulation models. Our results, using drugs of six different classes, demonstrate that exposure to sub-inhibitory antibiotic concentrations not only alters the dynamics of bacterial growth but also increases the mutation rate to antibiotic resistance and decreases the rate of production of persister cells thereby reducing the persistence level. Understanding this trade-off between mutation rates and persistence levels resulting from sub-inhibitory antibiotic exposure is crucial for optimizing, and mitigating the failure of, antibiotic therapy.

## INTRODUCTION

In the rational design of antibiotic therapy, drugs are administered such that the concentration of the treating drug exceeds the threshold needed to prevent the replication of the target pathogen [1]. However, in a treated individual, the concentration of an antibiotic within the body varies across different anatomical regions due to factors such as variations in vascularization and the pharmacokinetics (PK) of the treating antibiotic [2]. Notably, even though antibiotics are administered such that the concentration of the drug in the serum exceeds the minimum inhibitory concentration (MIC), they are often present at sub-inhibitory concentrations over time throughout the body [3]. Despite this, almost all studies on the pharmacodynamics (PD) of antibiotics focus on super-inhibitory concentrations, ignoring the effects of sub-inhibitory concentrations of antibiotics on bacteria.

In this study, we utilize a laboratory strain of the clinically significant pathogen *Staphylococcus aureus* [4] to examine the impact of exposure to sub-inhibitory concentrations of six antibiotic classes on growth dynamics, mutation rates, and the level of persistence. Persistence is the fraction of quiescent bacterial cells that survive treatment with a super-inhibitory concentration of an antibiotic [5]. In a previous study with *Escherichia coli*, we have demonstrated that exposure to sub-inhibitory concentrations of antibiotics results in a decrease in the growth rate along with the maximum bacterial density achieved, as well as an increase in the lag phase (the time before the bacterial population begins to replicate) [6]; we confirm the generality of those findings here. Moreover, other studies have established that super-inhibitory antibiotic concentrations can elevate the mutation rate for resistance to other drugs [7] we have found that this phenomenon extends to sub-inhibitory antibiotic concentrations as well. Finally, we provide evidence that pre-exposure to sub-inhibitory concentrations of antibiotics decreases the level of persistence.

## RESULTS

### The Effects of Sub-inhibitory Concentration of Antibiotics on Bacterial Growth Dynamics

To determine the effects of exposure to sub-inhibitory concentrations of antibiotics on the growth dynamics of bacteria, we follow the changes in the optical densities of *Staphylococcus aureus* Newman exposed to sub-inhibitory concentrations of antibiotics from six different classes in Fig. 1 [8]. The growth dynamics of *S. aureus* Newman vary among the drugs for all six antibiotics, however, there is a clear concentration-dependent variation in the maximum growth rate (Fig. S1), the maximum optical density (Fig. S2), and the lag time (Fig. S3). These results are consistent with those previously observed for *Escherichia coli* [6], demonstrating that the results obtained previously are not restricted to Gram-negative bacteria.

**Fig. 1.**
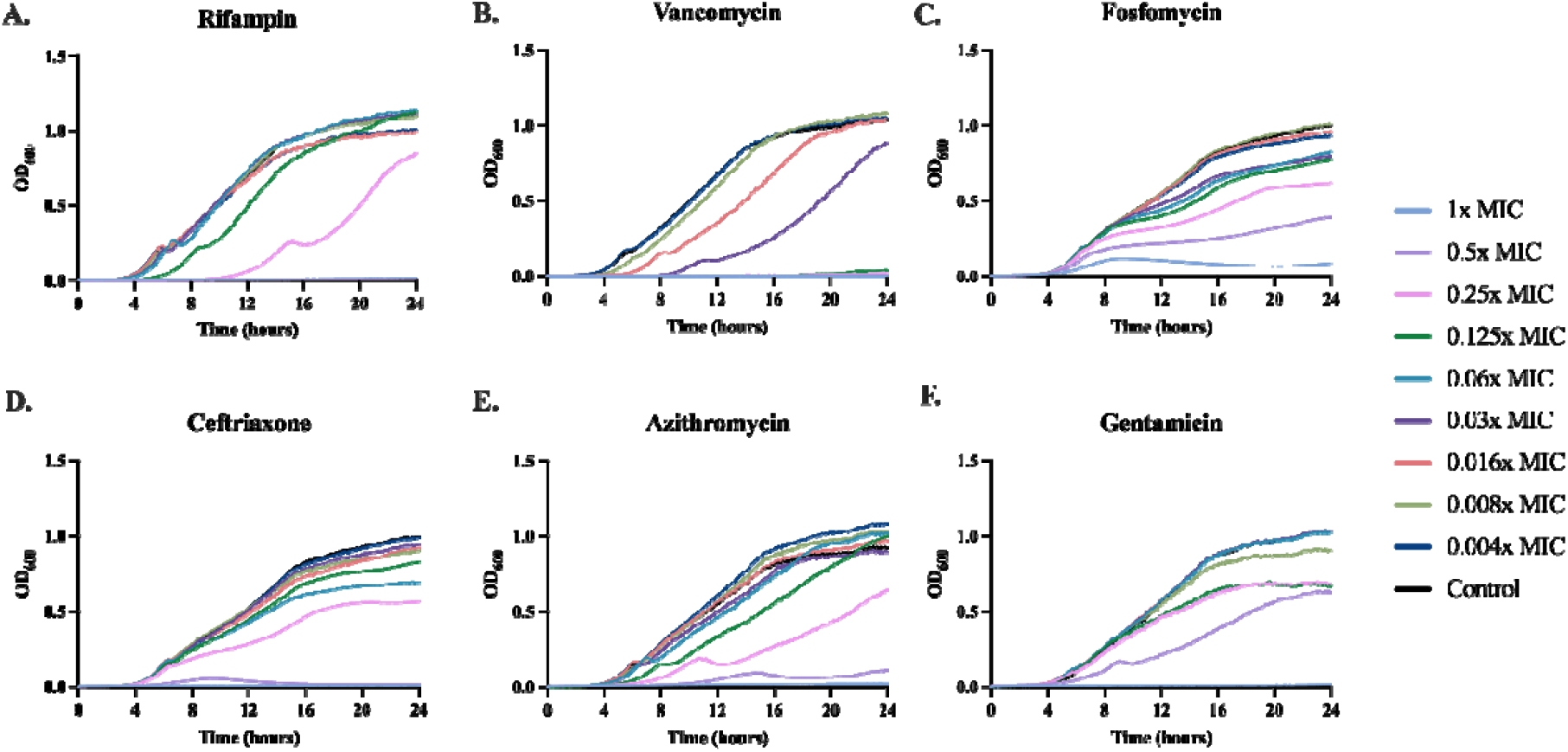
Growth dynamics of *S. aureus* with varying antibiotics and concentrations. Changes in the optical density at 600nm (OD600) exposed to different concentrations of six classes of drugs. Lines represent the average of five technical replicates. Each concentration is given as a fraction of the MIC shown in Table S1: 1x (light blue), 0.5x (light purple), 0.25x (pink), 0.125x (green), 0.06x (blue), 0.03x (purple), 0.016x (red), 0.008x (light green), 0.004x (dark blue), with a drug-free control shown in black.

### The Effects of Exposure to Sub-inhibitory Concentration of Antibiotics on the Mutation Rate

#### Null Model of Mutation Rate

To explore the intrinsic variation in the estimation of mutation rates, we use a mathematical-computer simulation model that employs the Monte Carlo process to generate mutants (Supplemental Text and Supplemental Equations 1-4) [9]. Shown in Table 1 are five independent runs of this model each with 20 independent replicates. Though there is variation in the estimated mutation rate between runs, this variation is not statistically significant.

**Table 1.** Variation in Mutation Rates Estimated from a Monte Carlo Simulation of Random Mutation.

|  | Null Model Mutation Rate Predictions |
| --- | --- |
| **Trial 1** | $2.96 \times 10^{-9}$ |
| **Trial 2** | $3.44 \times 10^{-9}$ |
| **Trial 3** | $3.43 \times 10^{-9}$ |
| **Trial 4** | $2.20 \times 10^{-9}$ |
| **Trial 5** | $2.98 \times 10^{-9}$ |

#### Changes in the Mutation Rate due to Sub-inhibitory Drug Pre-exposure

To determine the effect sub-inhibitory pre-exposure has on the mutation rate to antibiotic resistance, we exposed *S. aureus* Newman to the concentration of the six drugs above that did not change the maximum stationary phase density. After 24 hours of pre-exposure, we performed a Luria-Delbruck fluctuation test to determine the mutation rate to streptomycin resistance (Table 2) [10]. Notably, pre-exposure to sub-inhibitory concentrations of antibiotics significantly increased the mutation rate to streptomycin resistance, a result unanticipated by the null model.

**Table 2.**
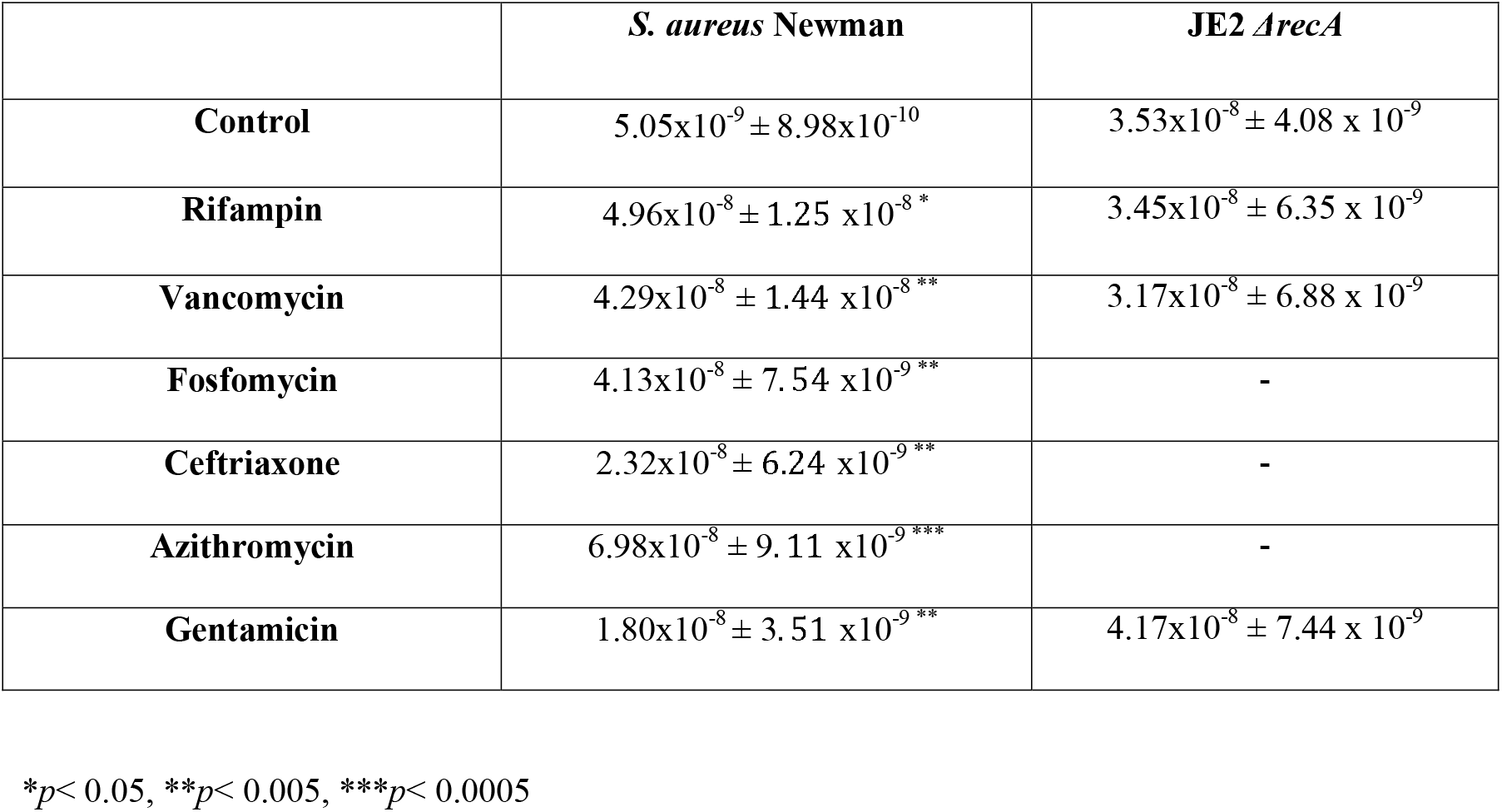
Mutation Rates to Streptomycin Resistance in *S. aureus* Pre-Exposed to Different Antibiotics.

|  | *S. aureus* Newman | JE2 *ΔrecA* |
| --- | --- | --- |
| **Control** | $5.05 \times 10^{-9} \pm 8.98 \times 10^{-10}$ | $3.53 \times 10^{-8} \pm 4.08 \times 10^{-9}$ |
| **Rifampin** | $4.96 \times 10^{-8} \pm 1.25 \times 10^{-8}^*$ | $3.45 \times 10^{-8} \pm 6.35 \times 10^{-9}$ |
| **Vancomycin** | $4.29 \times 10^{-8} \pm 1.44 \times 10^{-8}^{**}$ | $3.17 \times 10^{-8} \pm 6.88 \times 10^{-9}$ |
| **Fosfomycin** | $4.13 \times 10^{-8} \pm 7.54 \times 10^{-9}^{**}$ | - |
| **Ceftriaxone** | $2.32 \times 10^{-8} \pm 6.24 \times 10^{-9}^{**}$ | - |
| **Azithromycin** | $6.98 \times 10^{-8} \pm 9.11 \times 10^{-9}^{***}$ | - |
| **Gentamicin** | $1.80 \times 10^{-8} \pm 3.51 \times 10^{-9}^{**}$ | $4.17 \times 10^{-8} \pm 7.44 \times 10^{-9}$ |

\* $p < 0.05$ , \*\* $p < 0.005$ , \*\*\* $p < 0.0005$

To elucidate the contribution of the generalized bacterial stress response, known as the SOS response, to the increase in mutation rate, we repeated the above experiments with a strain lacking *recA*, the major constituent of the SOS response [11]. When this knockout strain was pre-exposed to the same fraction of the MIC of each drug, there was no evidence of a significant increase in the mutation rate (Table 2). The *recA* knockout was resistant to fosfomycin, ceftriaxone, and azithromycin; thus, these antibiotics could not be used for pre-exposure of this strain (Table S1). The background strain for this knockout, JE2, was found to have a higher baseline mutation rate than Newman (4.01×10^−8^ ± 8.87×10^−9^). However, when pre-exposed to sub-inhibitory concentrations of antibiotics, JE2 still exhibited a 10-fold increase in the mutation rate to streptomycin (p=0.008, n=20).

Streptomycin was the only drug used to estimate the mutation rate, although other antibiotics were tested. For this experiment, the mechanism of resistance must be a single point mutation, which significantly limits the classes of drugs that could be used. Tobramycin, another aminoglycoside, was found to have an extremely high baseline mutation rate (due to its inability to be enumerated on a fluctuation test), and thus any increase in the rate could not be observed. The fluoroquinolones were found to have too low of a baseline mutation rate, such that it was below the limit of detection. Interestingly, *S. aureus* Newman was found to be heteroresistant to the quinolone nalidixic acid, while it was not heteroresistant to the fluoroquinolone ciprofloxacin (Fig. S4).

### The Effects of Exposure to Sub-inhibitory Concentrations of Antibiotics on the Level of Persistence

#### Null Model of Persistence

To determine the effect that the rate of persistence generation has on the final level of persistence, we employed a mathematical-computer simulation model of persistence with differing rates of persister cell generation (Supplemental Text and Supplemental Equations 5-8). In Fig. 2, we show that a higher rate of persister cell generation results in a higher level of persistence at six hours, such that in a time-kill experiment the total number of surviving cells would be higher in a rate-dependent manner.

**Fig. 2.**
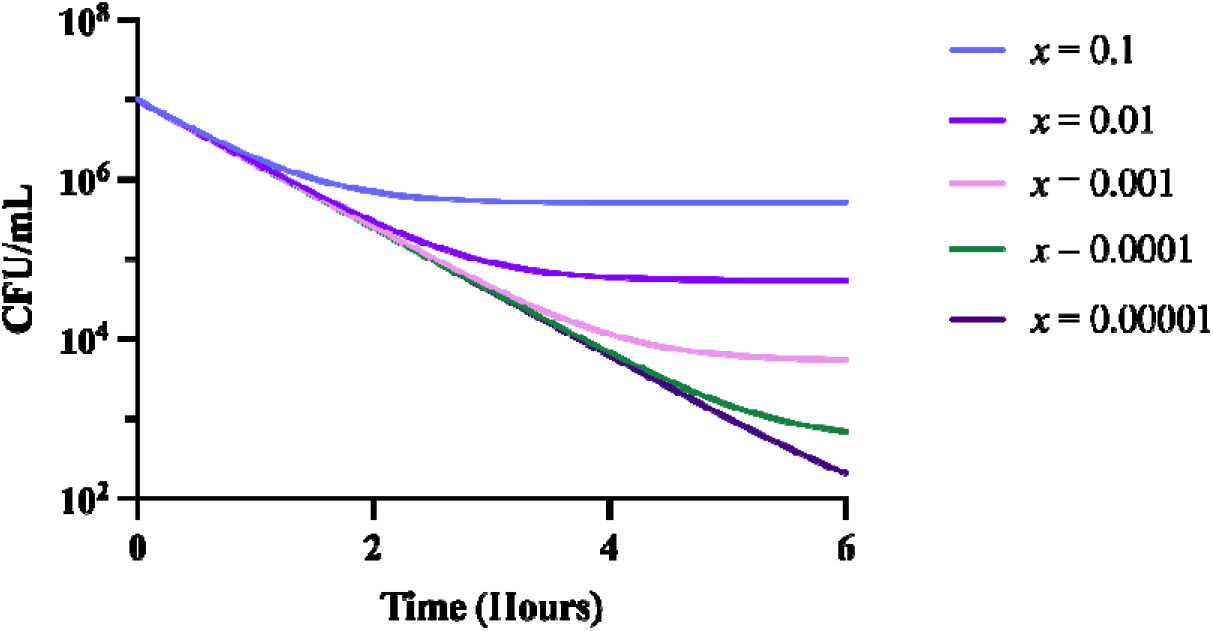
Predicted changes in the total cell density of a bacterial population capable of producing persister cells to a bactericidal antibiotic. These simulations assume all parameters are equal between runs except for the parameter *x*, the rate constant of persister cell generation. The other parameters used for this simulation are *A* = 5.0, *v*_S_ = 2.0, *v*_P_ = 0, *v*_MIN_ = -3.0, e = 5×10^−7^, *MIC* = 1.0, and *r* = 1000.

#### Changes in the Level of Persistence Due to Sub-inhibitory Drug Pre-exposure

To determine the effect that sub-inhibitory pre-exposure has on the level of persistence, we first had to select drugs for which *S. aureus* Newman shows persistence—which is shown on time-kill curves as cells that survive super-inhibitory drug exposure but do not replicate and do not have an increased MIC. In Fig. S5, we show that daptomycin and tobramycin, two highly bactericidal antibiotics, both have differing levels of persistence, whereas ciprofloxacin, tetracycline, and streptomycin do not exhibit clear evidence for persistence at the tested concentrations[12]. We chose 6x MIC for tobramycin and 4x MIC for daptomycin to perform subsequent time-kill curves to maximize the difference in the levels of persistence. To ensure the drug-exposed survivors were due to persistence and not some other phenomenon such as tolerance or resistance, single colonies from the last time point of the time-kills were selected, and the time-kill was repeated. The time-kill curves with these colonies were qualitatively and quantitatively similar to those in Fig. S5, showing that the surviving cells were indeed persisters (Fig. S6). MICs were performed on the cells surviving the time-kills and their MIC was found to be the same as the parental strain, providing evidence for persistence rather than resistance.

To elucidate the effects sub-inhibitory pre-exposure has on the level of persistence, we performed time-kill experiments with the drugs and concentrations selected above. Cultures were pre-exposed for 24 hours to the six antibiotics used in Fig. 1 at sub-inhibitory concentrations which were shown not to reduce the stationary phase densities. As shown in Fig. 3 and Fig. 4, pre-exposure to sub-inhibitory concentrations of the six antibiotics decreased the levels of persistence to both tobramycin and daptomycin. Variation in the initial density occurred due to the reduced densities generated by exposure to sub-inhibitory concentrations of the drugs.

**Fig. 3.**
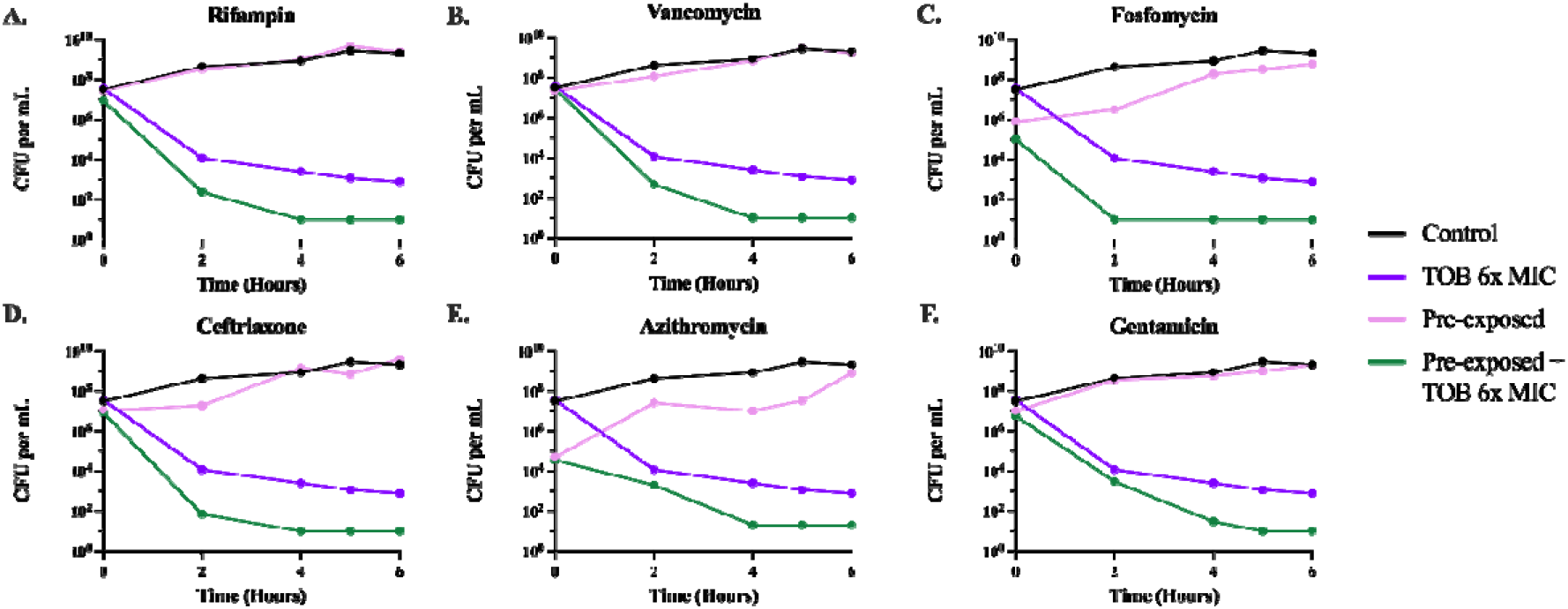
Time-kill Experiments with Tobramycin. Six-hour time-kill curves were performed with 6x the MIC of tobramycin (Table S1). Cultures were either pre-exposed for 24 h or not pre-exposed to sub-inhibitory concentrations of one of the six antibiotics; from there either the cultures were allowed to grow in the absence or presence of tobramycin. Lines represent: no pre-exposure, no tobramycin (black); no pre-exposure, tobramycin (purple); pre-exposure, no tobramycin (pink); and pre-exposure, tobramycin (green).

**Fig. 4.**
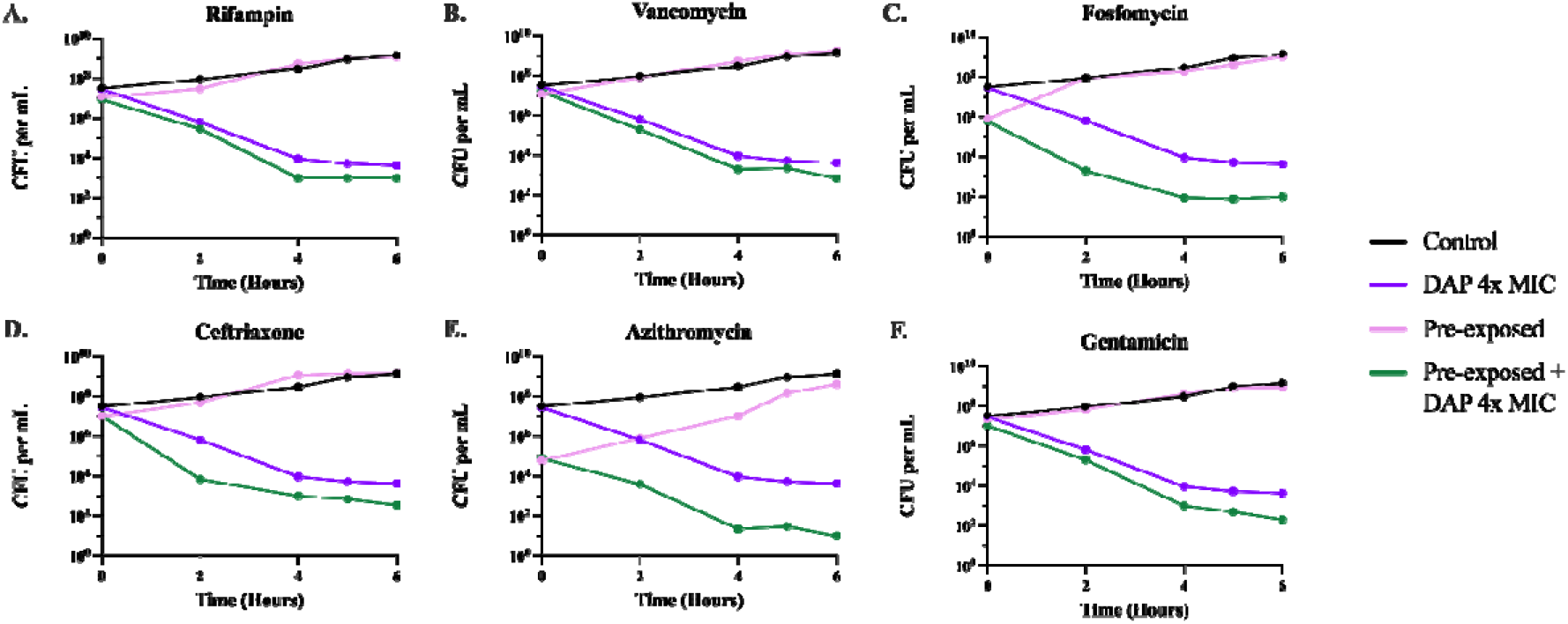
Time-kill Experiments with Daptomycin. Six-hour time-kill curves were performed with 4x the MIC of daptomycin (Table S1). Cultures were either pre-exposed for 24 h or not pre-exposed to sub-inhibitory concentrations of one of the six antibiotics; from there either the cultures were allowed to grow in the absence or presence of daptomycin. Lines represent: no pre-exposure, no daptomycin (black); no pre-exposure, daptomycin (purple); pre-exposure, no daptomycin (pink); and pre-exposure, daptomycin (green).

#### Changes in Metabolic Activity Due to Sub-inhibitory Drug Pre-exposure

Persister cells enter a state of dormancy in which they reduce their metabolic activity. Accordingly, if metabolism is increased, persistence levels will decrease [13]. To evaluate the effect that the pre-exposure to sub-inhibitory concentrations of antibiotics has on bacterial metabolic activity, we measured the intracellular amount of ATP via a luminescence assay. In Fig. 5 we show that pre-exposure to sub-inhibitory concentrations of the selected antibiotics increased the ATP levels, indicating a higher metabolic rate that may account for the results in Fig. 3 and Fig. 4.

**Fig. 5.**
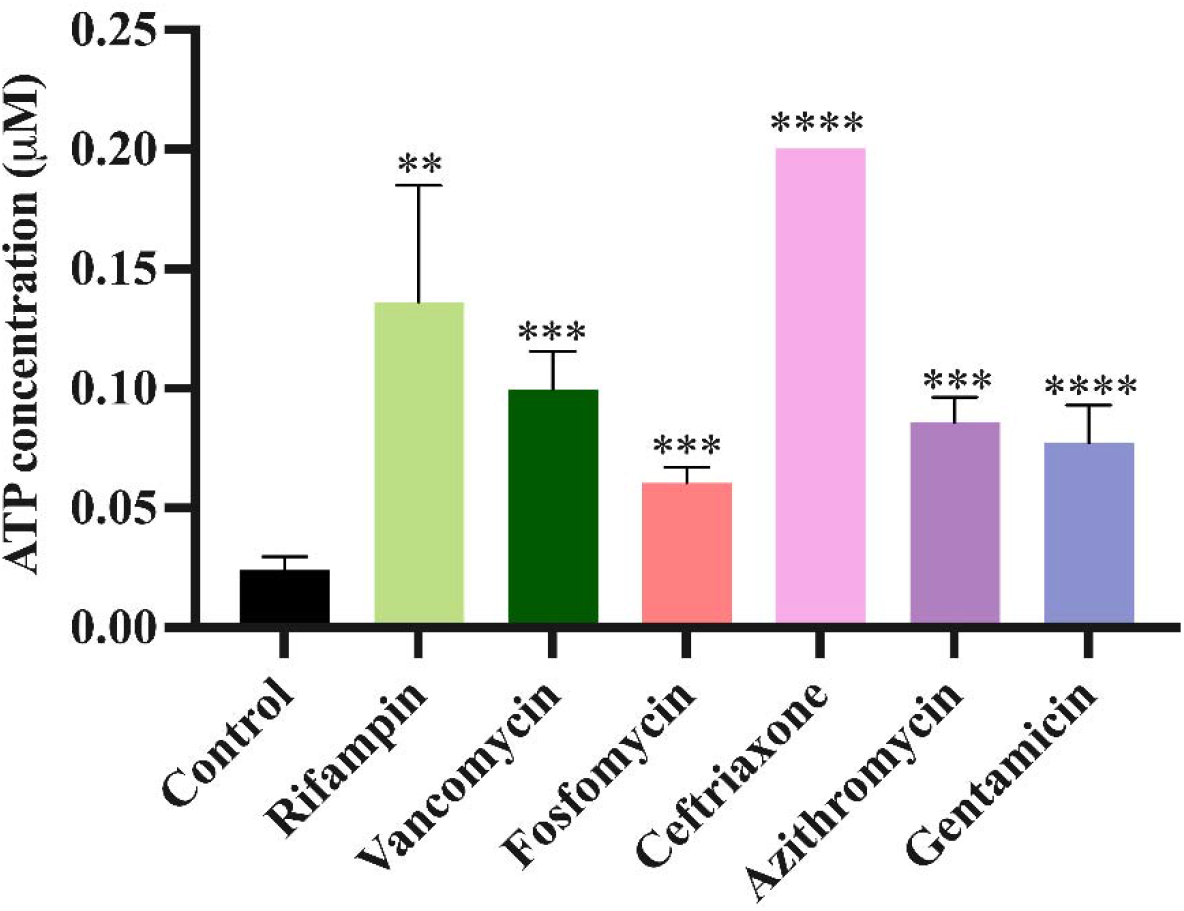
ATP determination. Cultures were either pre-exposed or not pre-exposed to sub-inhibitory concentrations of one of the six antibiotics: From there the amount of ATP in these cultures was experimentally estimated after 24 hours of pre-exposure via luminescence at 560 nm. \*\**p*< 0.005, \*\*\**p*< 0.0005, \*\*\*\**p*< 0.00005.

## DISCUSSION

Antibiotics are prescribed to patients at concentrations designed to exceed the minimum concentration necessary to prevent the replication of the target pathogen [14]. Therefore, the minimum inhibitory concentration (MIC) is the dominant and often the unique pharmacodynamic parameter used to design antibiotic treatments [15]. However, i*n vivo* conditions introduce significant variability in factors such as local bacterial concentration at the infection site, replication rate, nutrient availability, and the immune response [16]. Moreover, though the antibiotic is administrated at super-inhibitory concentrations, this concentration may not be reached in all, or even most, locations of the body, including the infection sites [17]. This means treatment occurs at gradients of antibiotic concentrations throughout the body, including antibiotic concentrations insufficient to kill or prevent the replication of the infecting bacteria [18].

Previous studies have revealed that exposing *E. coli* to sub-inhibitory concentrations of antibiotics leads to decreasing both maximum growth rate and maximum optical density while increasing the lag phase of growth [6]. Our results here confirm this phenomenon applies to *S. aureus* as well. These changes are consistent through all six classes of drugs tested where a concentration-dependent response is observed; as the concentration of the antibiotic increases, so does the degree of impairment of the growth dynamics. These results show that significant antibacterial activity occurs at sub-inhibitory concentrations, in some cases exceptionally lower than the MIC, suggesting that antibiotic may have clinical utility at sub-inhibitory concentrations. This may explain why infections can be successfully treated despite being located in sites where super-inhibitory antibiotic concentrations are not achieved. Apart from locational heterogeneity, sub-inhibitory antibiotic concentrations can also obtain due to suboptimal dosing, extending the time between doses, and using partially inactivated drugs due to inappropriate storage.

Along with the changes in growth dynamics, sub-inhibitory exposure may lead to physiological changes in the bacteria [19]. When bacteria are exposed to super-inhibitory concentrations of antibiotics, resistant mutants in the population will be able to survive and replicate in the presence of this selective pressure due to mutations [20, 21]. Mutation rates, including those of antibiotic resistance, are not fixed. One pathway that modulates mutation rates is the SOS response which is nearly ubiquitous in bacteria [22]. This response plays a vital role in DNA repair and enables survival under physiological stress. Several external factors can lead to the activation of the SOS response [23]. Our results illustrate one of these factors is exposure to sub-inhibitory concentrations of antibiotics (Table 2). The major regulator of the SOS response is RecA [11]. In *S. aureus*, there are two major pathways involved in this response: the LexA dependent pathway which results in the expression of the UmuC error prone polymerase, and the RexAB dependent pathway which results in the formation of small colony variants [24-26]. Taken together, activation of both pathways results in an increase in the mutation rate of at least one order of magnitude, as we have shown in Table 2. As expected, when *recA* is knocked out, these pathways cannot be activated, and pre-exposure to sub-inhibitory concentrations of antibiotics does not lead to a change in the mutation rate.

Another phenomenon that could arise from pre-exposure to sub-inhibitory concentrations of antibiotics is changes in persistence levels. Persistence is a temporary phenotypic change in which the majority of the population is susceptible to antibiotics and a minority population is capable of surviving exposure to antibiotics without an increase in the MIC [27, 28]. Persister cells can survive antibiotic treatment by entering a dormant or slow-growing state, due to several possible mechanisms [29, 30]. Different environmental factors can change the frequency of generation of persister cells in a bacterial population; our results here show that one of these factors is the exposure to antibiotics—in this case, sub-inhibitory levels of six distinct antibiotics. When bacteria are confronted with sub-inhibitory levels of antibiotics before encountering super-inhibitory concentrations of other drugs, it triggers metabolic changes which decrease the rate of generation of these persister cells. Our results further demonstrate that these metabolic changes occur due to exposure to sub-inhibitory concentrations of antibiotics which is shown by a higher intracellular ATP concentration (Fig. 5). This increase in metabolic activity opposes the dormancy that defines persistence, therefore leading to a lower rate of persister cell formation when the bacterial populations are then exposed to super-inhibitory concentrations of other drugs. Unexplored, but testable, implications also arise from this increase in metabolic activity. Conceivably, toxins and other virulence factors are also upregulated by exposure to sub-inhibitory concentrations of antibiotics.

These results contribute to our understanding of the interaction between bacterial mutation, persistence, and antibiotics as an academic matter; however, there are serious clinical implications that follow these findings as well [31]. The administration of a first line of antibiotic therapy will create a gradient of antibiotic concentrations within the body. If this first treatment fails, and a secondary line of treatment is administered, the increase in mutation rate produced in response to the sub-inhibitory concentrations in different body locations could lead to the generation of resistant mutants which could then result in treatment failure that would not otherwise have occurred. On the other hand, we show that persistence would be reduced wherever there was pre-exposure to antibiotics. This ability to persist is an important attribute for bacterial populations when conditions are unfavorable for their survival. As a result, pre-exposure to sub-inhibitory concentrations of antibiotics reducing persistence levels could enhance second-line treatment efficacy, improving the effectiveness of super-inhibitory concentrations of the antibiotic used in therapy and therefore reducing the risk of recurrent infections [32]. These results are especially salient in chronic and recurrent infections such as those involving biofilms [33]. Ultimately, these findings boil down to one important trade-off that has real-world impacts in the clinic, that is a trade-off between higher mutation rates and lower persistence levels resulting from previous exposure to sub-inhibitory concentrations of antibiotics.

## MATERIALS AND METHODS

### Growth media

All experiments were conducted in Muller Hinton II (MHII) Broth (90922-500G) obtained from Millipore. All bacterial quantification was done on Lysogeny Broth (LB) agar (244510) plates obtained from BD. E-tests were performed on MH agar plates made from MH broth (M391-500g) with 1.6% agar obtained from HiMedia.

### Growth Conditions

Unless otherwise stated, all experiments were conducted at 37°C with shaking.

### Bacterial strains

All experiments were performed with *Staphylococcus aureus* Newman obtained from Bill Schafer of Emory University. Je2Δ*recA* and Je2 from the Nebraska Transposon Mutant Library [34] were obtained from Joanna Goldberg of Emory University.

### Antibiotics

Streptomycin (S6501), sulfamethoxazole (S6377), vancomycin (V1130), ceftriaxone (C5793), fosfomycin (P5396), and daptomycin (D2446) were all obtained from Sigma-Aldrich. Tobramycin (T1598) was obtained from Spectrum. Azithromycin (3771) was obtained from TOCRIS. Ciprofloxacin (A4556) was obtained from AppliChem Panreac. Gentamicin (BP918-1) and rifampin (BP2679-1) were obtained from Fisher. Nalidixic acid (KCN23100) was obtained from PR1MA. Tetracycline (T17000) was obtained from Research Products International. All E-test strips were obtained from Biomérieux.

### Sampling bacterial densities

The densities of bacteria were estimated by serial dilution in 0.85% saline and the total density of bacteria was estimated on LB plates with 1.6% agar.

### Growth rate estimation

Exponential growth rates were estimated from changes in optical density (OD600) in a Bioscreen C For this, 24-hour overnight cultures were diluted in MHII to an initial density of approximately 10^5^ cells per mL. Five technical replicates were performed for each condition in a 100-well plate. The plates were incubated at 37°C and shaken continuously. Estimates of the OD600 were made every 5 minutes for 24□hours. Normalization was performed and then means and standard deviations of the maximum growth rate, lag time, and maximum OD were found using an R Bioscreen C analysis tool accessible at https://josheclf.shinyapps.io/bioscreen_app.

### Minimum inhibitory concentration estimation via broth microdilution

MICs were determined according to the CLSI guidelines, deviating only in the choice of media [35]. Briefly, 96-well plates with two-fold dilutions of antibiotics in MHII media were prepared and inoculated with 10^5^ bacteria per mL. An extended gradient was created by combining three sets of two-fold serial dilutions from three starting antibiotic concentrations. The plates were incubated at 37°C with conditions shaking and the optical density (OD600) was measured after 24 hours.

### Fluctuation tests

Independent overnights of *S. aureus* Newman, Je2Δ*recA*, and Je2 were either exposed to sub-inhibitory concentrations of rifampin at 0.5x MIC, vancomycin at 0.5x MIC, fosfomycin at 0.25x MIC, ceftriaxone at 0.25x, gentamicin at 0.25x, azithromycin at 0.25x, or grown without antibiotic and then plated on LB agar plates containing 5x MIC of streptomycin. Experiments were performed with 20 biological replicates and the mutation rates were calculated as in [36, 37] with BZrates.com.

### Time kill experiments

Cultures of 10^7^ *S. aureus* Newman were either exposed overnight to sub-inhibitory concentrations of rifampin, vancomycin, fosfomycin, ceftriaxone, azithromycin, and gentamicin at the above concentrations or were grown without antibiotics. After this overnight incubation, all cultures were diluted in fresh MHII to 10^7^ cells per mL. The cultures were then exposed to super-MIC concentrations of streptomycin, daptomycin, tetracycline, tobramycin, or ciprofloxacin at varying concentrations, and viable cell density was estimated at 0, 2, 4, 5, and 6 hours.

### Population analysis profile test

PAP tests were performed as in [38, 39]. Briefly, a gradient of nalidixic acid or ciprofloxacin concentrations was added to LB plates. The concentrations were 0, 0.5, 1, 2, 4, 8, 16, and 32 xMIC. Multiple dilutions of *S. aureus* Newman (10^0^-10^−7^) were then plated on every concentration. Colonies were enumerated after 48 hours and the highest dilution with colonies present was recorded. The frequency of surviving cells was calculated by dividing the highest density of cells at each concentration by the number of surviving cells on plates with no antibiotics.

### Numerical solutions (simulations)

For our numerical analysis of the mathematical models detailed in the Supplemental Text, we used Berkeley Madonna, using parameters in the ranges estimated for *S. aureus* Newman. Copies of the Berkeley Madonna program used for these simulations are available at www.eclf.net.

### Statistical Analysis

Statistical significance analysis was carried out by paired t-tests using GraphPad Prism (version 10.2.0).

### ATP Assay

ATP determination kits were obtained from ThermoFisher Scientific (A22066). To perform the ATP determination, the manufacture’s provided protocol was followed with the following changes. Overnight cultures either pre-exposed to the antibiotics or not exposed were pelleted and the pellets washed with saline. Cultures were resuspended in saline and sonicated with a Branson Needle-Tip Sonicator. Post-sonication, cells were centrifuged, and the supernatants were placed in a black 96-well plate and incubated at room temperature for 30 minutes. After incubation, luminescence was then read at 560 nm.

## Supporting information

All Supplemental Text

## Acknowledgments

We thank the U.S. National Institute of General Medical Sciences for their funding support via R35 GM 136407 and the U.S. National Institute of Allergy and Infectious Diseases for their funding support via U19 AI 158080. FB acknowledges the support of CIBERESP (CB06/02/0053) from the Carlos III Institute of Health of Spain. The funding sources had no role in the design of this study and will not have any role during its execution, analysis, interpretation of the data, or drafting of this report. The content is solely the responsibility of the authors and do not necessarily represent the official views of the National Institutes of Health nor those of the Carlos III Institute of Health of Spain.

## Author Contributions

**Conceptualization:** ASI, BAB, TGG, JAM, APS, BRL

**Methodology:** ASI, BAB, TGG, BRL

**Investigation:** ASI

**Visualization:** ASI, TGG

**Funding Acquisition:** BRL

**Project Administration:** BRL

**Supervision:** BRL

**Writing– Initial Draft:** ASI, BAB, TGG, APS, FB, BRL

**Writing– Review & Editing:** ASI, BAB, TGG, JAM, APS, FB, BRL

**Competing Interest Statement:** The authors have no competing interests to declare.

## Notes

### Competing Interest Statement

The authors have declared no competing interest.

