## Supplementary material for "The Tradeoffs Between Persistence and Mutation Rates at Sub-Inhibitory Antibiotic Concentrations in *Staphylococcus aureus*": All Supplemental Text

**This PDF file includes:**

Supplemental Text

Supplemental Equations 1 to 8

Supplemental Figures 1 to 6

Supplemental Table 1

Supplemental References

**SUPPLEMENTAL TEXT**

**Fluctuation Test Simulations**

There are two populations of bacteria, *Y* and *Z*, with densities (cells per mL) at a single limiting resource, *r* (µg/mL). *Y* is the ancestral population and *Z* is the population of mutants generated from *Y*. The *Y* and *Z* populations grow at maximum rates, *v_Y_* and *v_Z_* per cell per hour, respectively. The respective net growth rates are equal to the product of the resource function, *ψ(r)* (Equation 1), and the maximum growth rate (Equations 2 and 3). The parameter *k* (µg/mL) is the concentration of the limiting resource where the rate of growth of the population is half its maximum value. The limiting resource is consumed at a rate proportional to the product of the sum of the maximum growth rates and densities of the populations, the resource function, *ψ(r),* and the conversion efficiency parameter, *e* (µg/mL), which is the amount of the limiting resource required to produce a new cell (Equation 4) [1, 2].

With these definitions and assumptions, the rates of change in the densities of the *Y* and *Z* populations, as well as the concentration of the limiting resource, are given in the below set of equations. 1/(*VOL***dt*) is the number of mutants generated by a Monte Carlo process [3] during the finite interval *dt* hours. In this Monte Carlo process, using the Euler Method, a pseudorandom number x (0 ≤ x ≤1) is generated from a rectangular distribution at every step, *t*+*dt*, where *dt* is the step size. The parameter *Q* can take two values, either 0 or 1. Given the parameter *µ* (the mutation rate per cell per hour) if x < *µ*$\cdot Y\cdot\psi(r)\cdot dt\cdot VOL$ then *Q* is 1. If x ≥ *µ*$\cdot Y\cdot\psi(r)\cdot dt\cdot VOL$ then *Q* is 0.

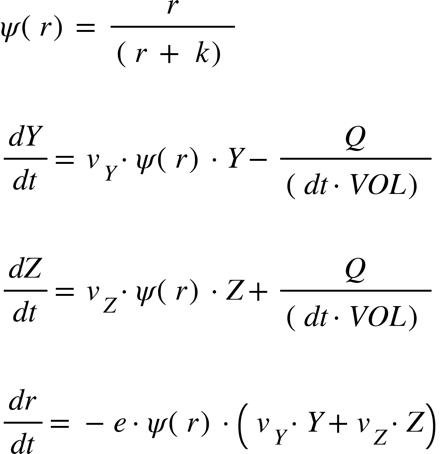

Equation 1

Equation 2

Equation 3

Equation 4

**Model of Bacterial Growth in the Presence of an Antibiotic with Persistence**

There are two populations of bacteria susceptible and persisters with designations and densities, *S* and *P* (cells per mL), a limiting resource of concentration *r* (µg/mL), and an antibiotic *A* (µg/mL). The growth of the *S* population is limited by both the concentration of the resource and the antibiotic, while the growth of the *P* population is limited only by the concentration of the resource (Equation 1). The *P* population is generated directly via transition from the *S* population at the rate *x* (per cell per hour). The *S* and *P* populations grow at a maximum rate *v_S_* and *v_P_* (per cell per hour, respectively). The net growth rate of the *S* population is a function of the concentration of the antibiotic, the maximum growth rate, the minimum growth rate (which is less than zero), the MIC, and the parameter *κ* (Equation 5) [4]. *MIC* is the minimum inhibitory concentration of the antibiotic. *κ* is a measure of the steepness of the function; the greater the value of *κ*, the more acute the steepness.

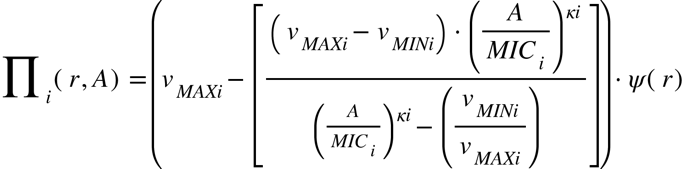

Equation 5

With these definitions and assumptions, the rate of change in the density of the different bacterial populations and changes in the concentration of the limiting resource are given by the set of coupled differential equations in Supplemental Equations 6-8.

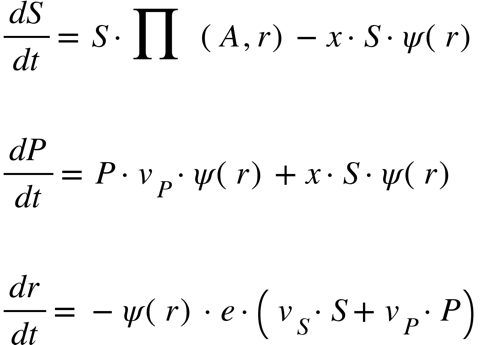

Equation 6

Equation 7

Equation 8

**SUPPLEMENTAL FIGURES**
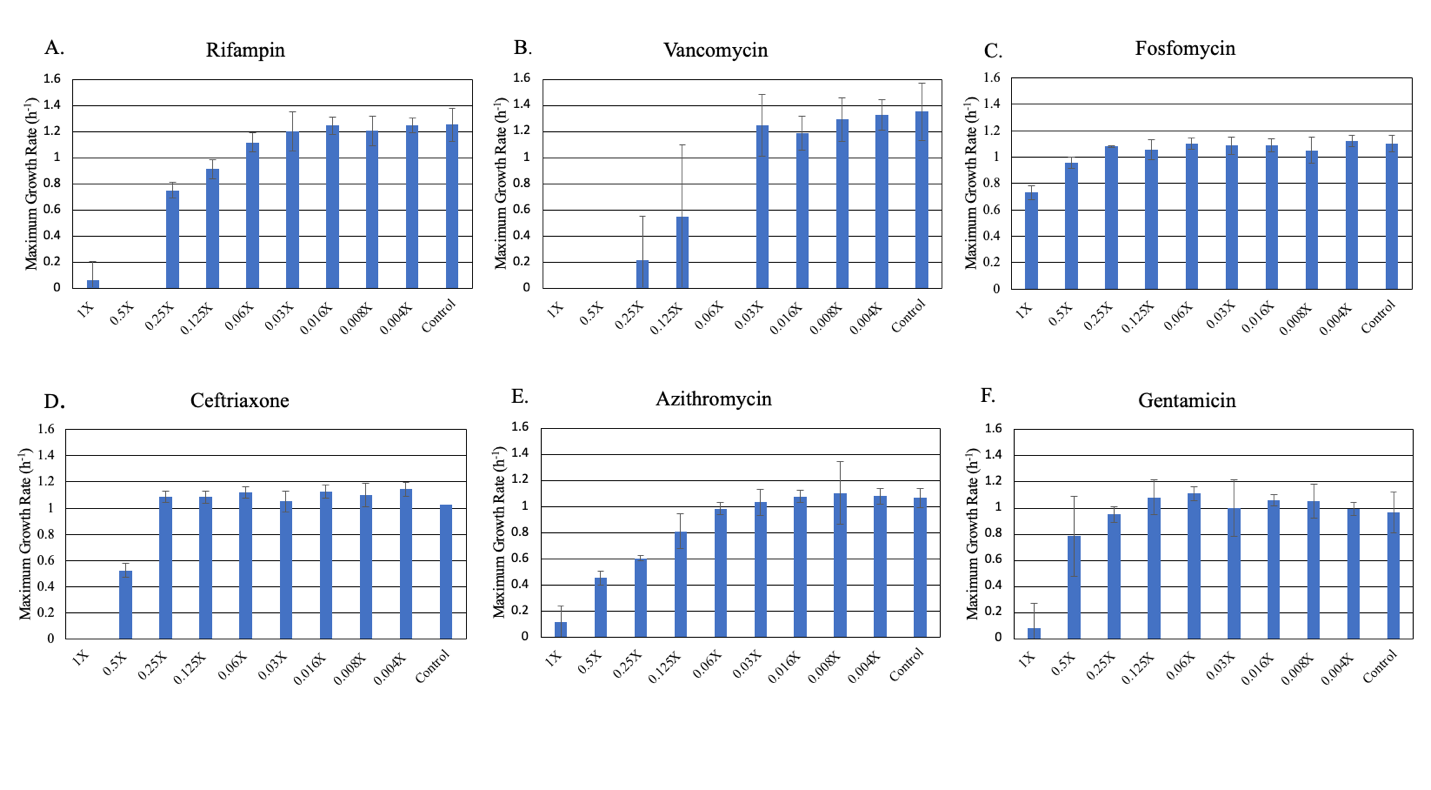

**Fig. S1. Changes in maximum growth rate (v_MAX_)** **of *S. aureus* Newman exposed to different sub-inhibitory concentrations of six antibiotics for 24 hours in MHII**. Bars are representative of the average of five technical replicates. Each concentration is shown as a fraction of the MIC for the noted drug: (A) rifampin, (B) vancomycin, (C) fosfomycin, (D) ceftriaxone, (E) azithromycin, (F) gentamicin.

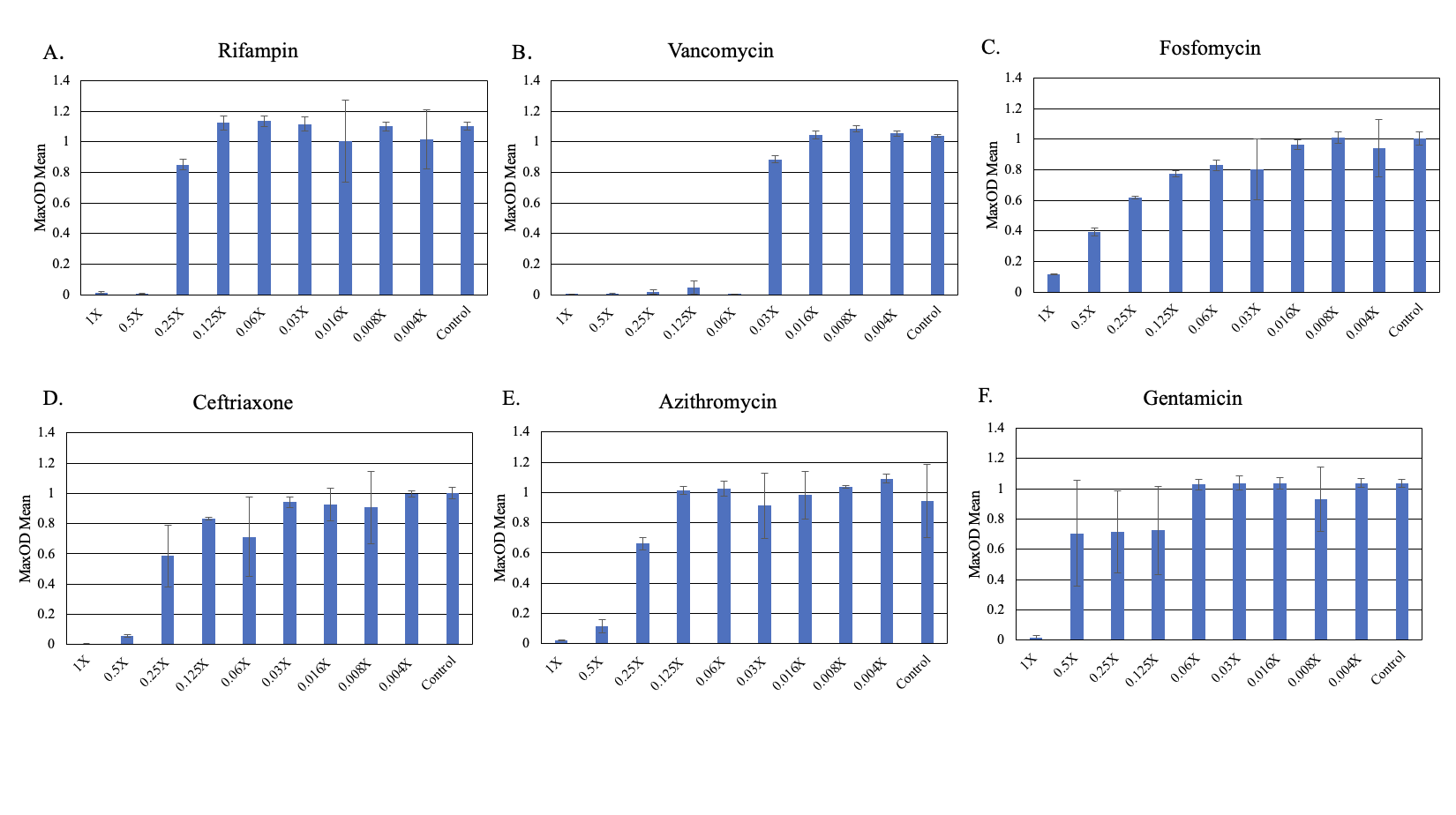

**Fig. S2. Changes in the maximum optical density (OD 600nm)** ***S. aureus* Newman exposed to different sub-inhibitory concentrations of six antibiotics for 24 hours in MHII**. Bars are representative of the average of five technical replicates. Each concentration is shown as a fraction of the MIC for the noted drug: (A) rifampin, (B) vancomycin, (C) fosfomycin, (D) ceftriaxone, (E) azithromycin, (F) gentamicin.

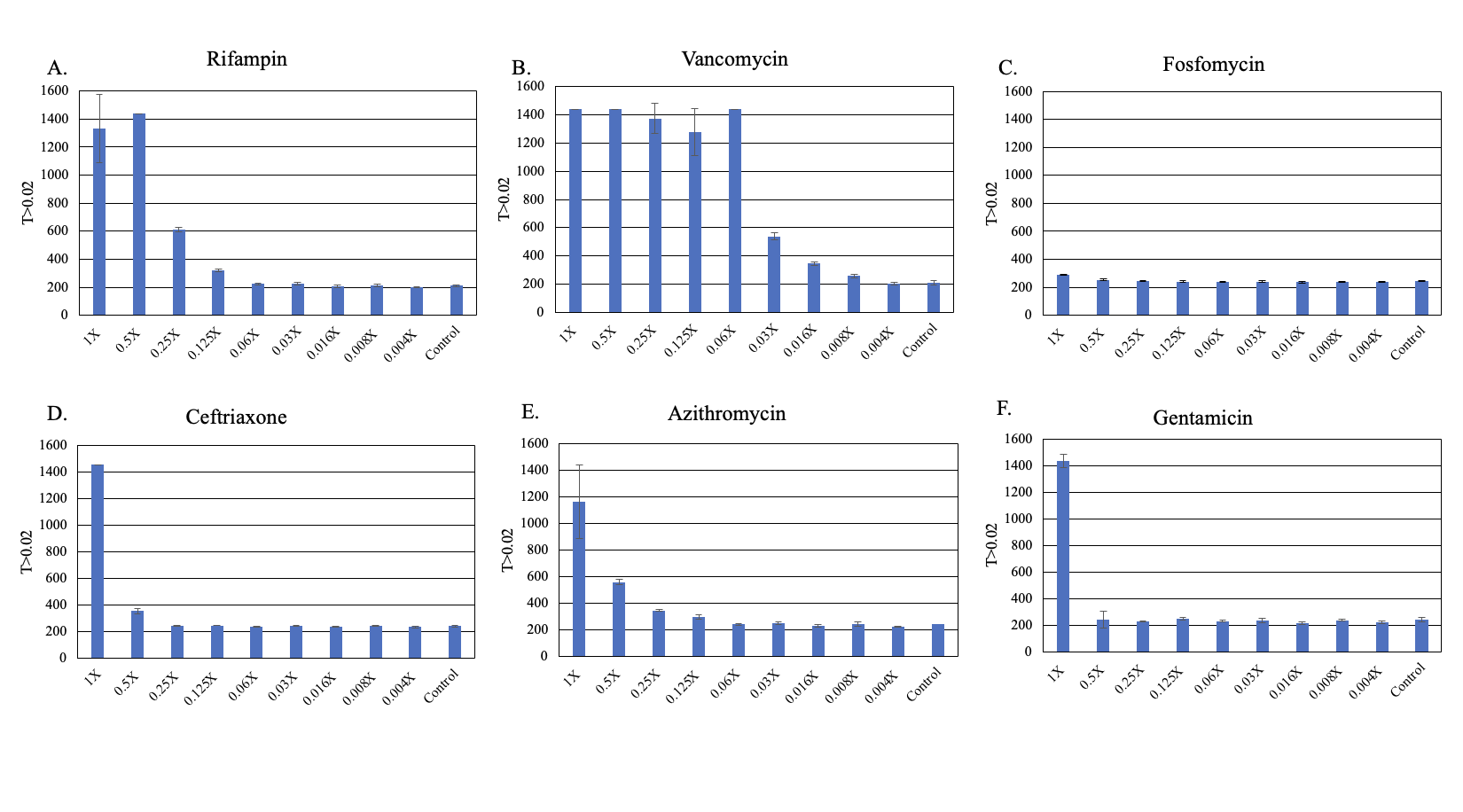

**Fig. S3. Changes in the time before the bacteria start to grow (lag time)** ***S. aureus* Newman exposed to different sub-inhibitory concentrations of six antibiotics for 24 hours in MHII**. Bars are representative of the average of five technical replicates. Each concentration is shown as a fraction of the MIC for the noted drug: (A) rifampin, (B) vancomycin, (C) fosfomycin, (D) ceftriaxone, (E) azithromycin, (F) gentamicin.

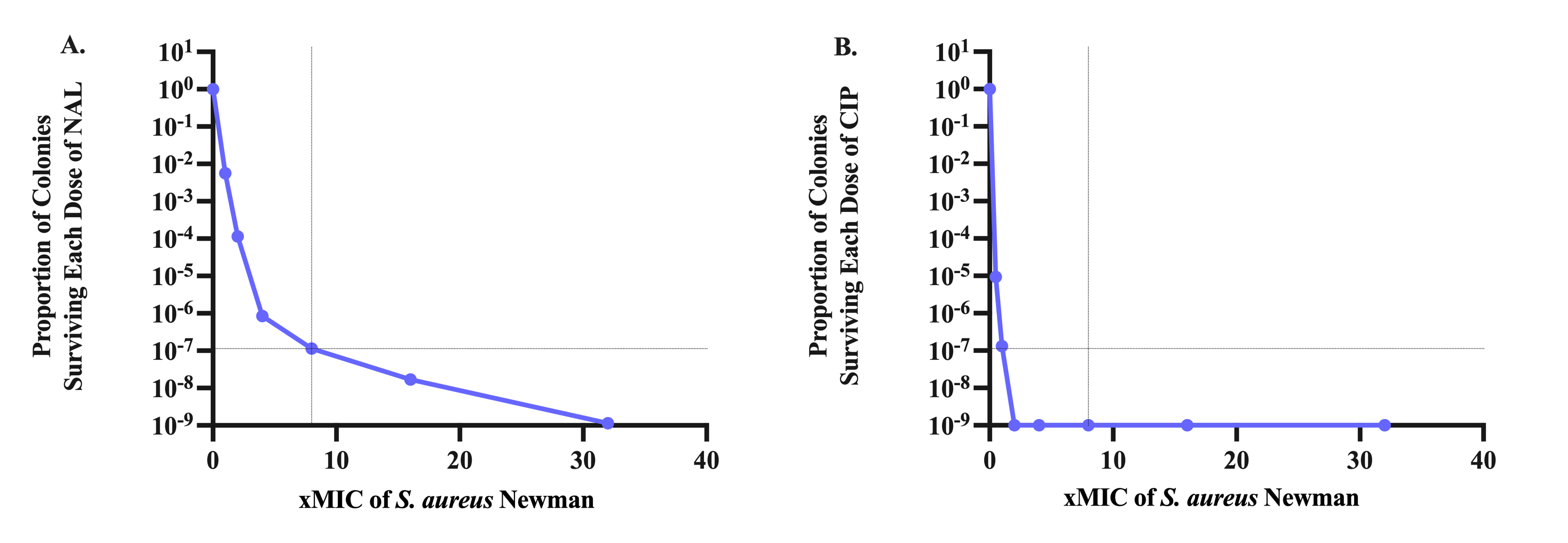

**Fig. S4. Population Analysis Profile (PAP) test of nalidixic acid and ciprofloxacin.** The ratio of the density of the number of bacteria surviving at an antibiotic concentration relative to that surviving in the absence of the antibiotic. (A) Nalidixic acid, (B) ciprofloxacin.

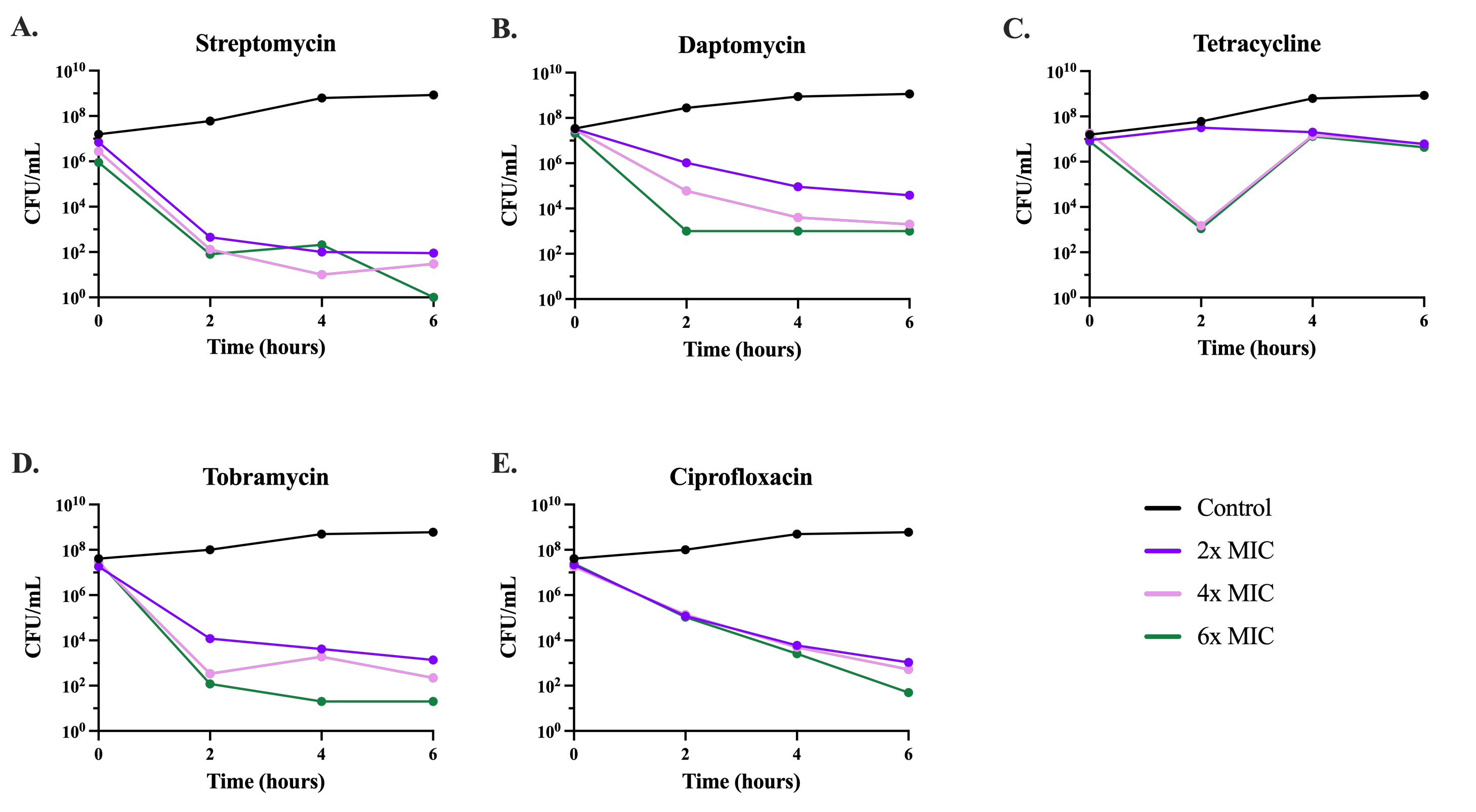

**Fig. S5. Time-kill experiments determining the level of persistence to different antibiotics.** Six-hour time-kill experiments were performed with (A) streptomycin, (B) daptomycin, (C) tetracycline, (D) tobramycin, and (E) ciprofloxacin at differing concentrations. Lines represent no selected drug (black); 2x the MIC of the selected drug (purple); 4x the MIC of the selected drug (pink); and 6x the MIC of the selected drug (green).

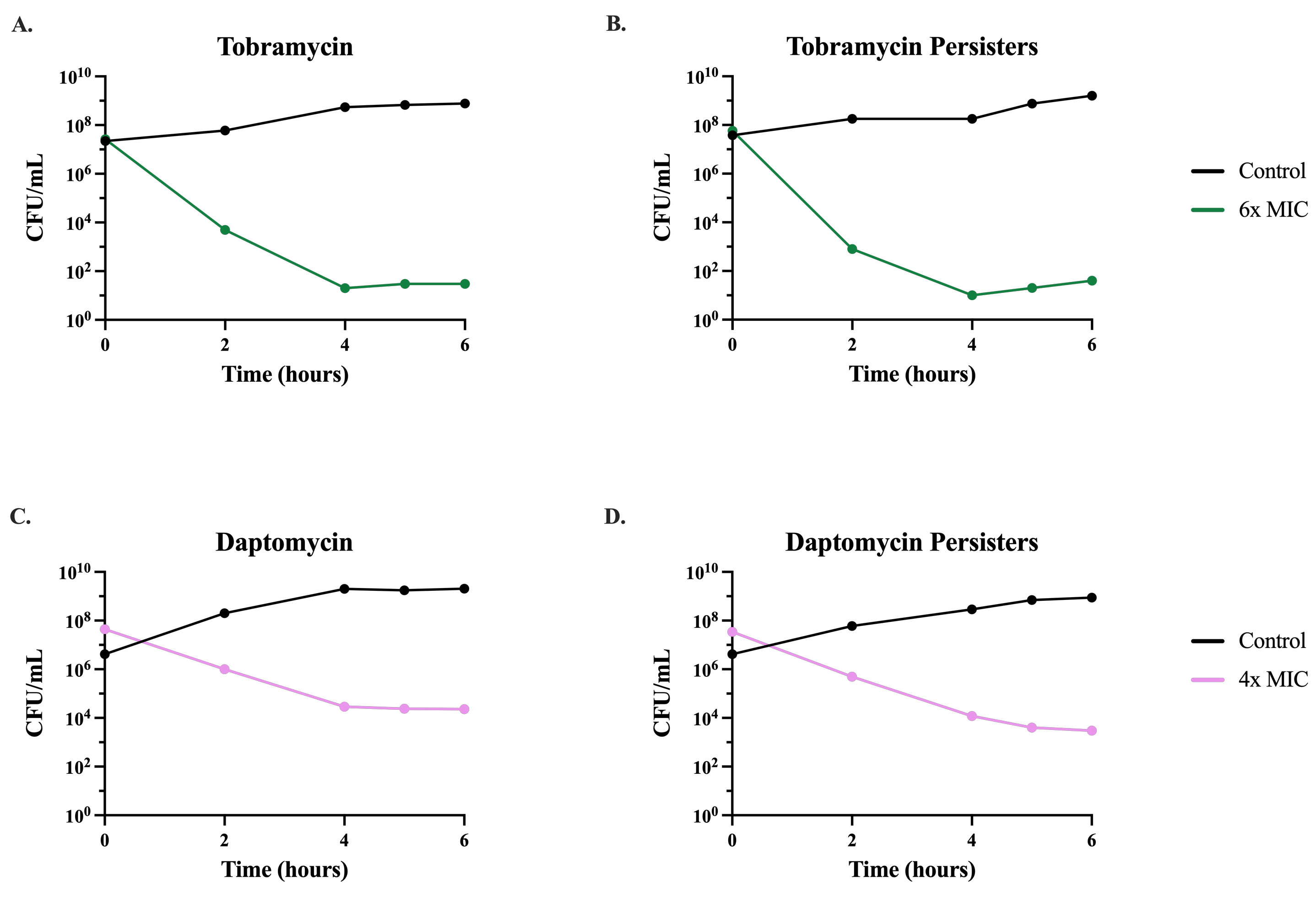

**Fig. S6. Time-kill experiments confirming persistence.** Six-hour time-kill experiments were performed with tobramycin (A and B) and daptomycin (C and D) at 6x and 4x respectively. Time kills checking for persisters (B and D) were performed with colonies selected from the previous time kills (A and C). Lines represent no drug (black); 4x the MIC of the selected drug (pink); and 6x the MIC of the selected drug (green).

**Table S1.** **Experimentally estimated MICs of *S. aureus* Newman for thirteen antibiotics in MHII broth.**

| **Antibiotic** | ***S. aureus* Newman**  **MIC** (µg/mL) | **Je2 *ΔrecA* MIC** (µg/mL) | **Je2 MIC** (µg/mL) |
| --- | --- | --- | --- |
| Rifampin | 0.004 | 0.012 | >1024 |
| Vancomycin | 2 | 2 | 2 |
| Fosfomycin | 2 | >1024 | >1024 |
| Ceftriaxone | 4 | >1024 | >1024 |
| Azithromycin | 2 | >1024 | >1024 |
| Gentamicin | 1 | 1 | >1024 |
| Nalidixic Acid | 16 | - | - |
| Ciprofloxacin | 0.50 | - | - |
| Tobramycin | 0.50 | - | - |
| Streptomycin | 12 | - | - |
| Daptomycin | 2 | - | - |
| Tetracycline | 1.50 | - | - |
| Sulfamethoxazole | >1024 | - | - |
